# Extracellular electron transport in coral-algal symbiosis studied by chlorophyll fluorescence relaxation and ferricyanide reduction

**DOI:** 10.1101/2025.11.26.690407

**Authors:** Sabit Mohammad Aslam, Priyanka Pradeep Patil, Szilárd Kovács, Anthony W. D. Larkum, Milán Szabó, Imre Vass

**Author notes:** Corresponding authors: Imre Vass,; Milán Szabó.

## Abstract

Coral health depends on intricate metabolic interactions between the coral host and its symbiotic algae, Symbiodiniaceae. While nutrient exchange is well established, electron-level interactions have remained unexplored. Here, we provide evidence for extracellular electron transport (EET) within coral-algal symbiosis, supported by variable chlorophyll (Chl) fluorescence and ferricyanide reduction measurements. We observed a slow wave in the relaxation of flash-induced Chl fluorescence kinetics under microaerobic conditions in both isolated *Fugacium kawagutii* (CS156) cells and intact corals, reflecting redox dynamics of the primary quinone electron acceptor (Q_A_) in the photosynthetic electron transport chain. The addition of the extracellular electron acceptor ferricyanide decreased the wave amplitude and Q_A_ reduction while being reduced to ferrocyanide, demonstrating EET from the symbiont to extracellular acceptors. Slower Chl fluorescence rise kinetics under continuous illumination in intact corals compared to isolated symbionts indicate that electrons may also flow from symbionts to the host. Under low oxygen conditions, Q_A_ was gradually reduced in corals in darkness but not in isolated symbiont cultures, suggesting electron transfer from host to symbiont. Together, these results indicate bidirectional extracellular electron exchange between symbiotic partners, pointing to a previously unrecognized mechanism for redox balancing in coral-algal symbiosis. This pathway likely contributes to metabolic resilience and the maintenance of coral health under fluctuating environmental conditions.

## Introduction

Coral reefs are among the most complex and diverse ecosystems on Earth, despite existing in nutrient-poor marine environments. Covering less than 0.1% of the ocean floor, they support approximately 25% of all marine species, making them indispensable to marine biodiversity. Beyond their ecological significance, coral reefs play a crucial role in the global carbon cycle by acting as carbon sinks, absorbing carbon dioxide from the atmosphere. Their ability to thrive in nutrient-deficient waters is largely due to a mutualistic relationship with unicellular photosynthetic algae known as zooxanthellae^1^. These algae reside within coral tissues and drive primary production by converting the energy of sunlight into chemical energy through photosynthesis. This process not only fuels coral metabolism but also facilitates nutrient recycling within the reef ecosystem, sustaining an otherwise energy-limited environment. Over time, the photosynthetic machinery of these dinoflagellate symbionts has developed specialized mechanisms to optimize light energy utilization and prevent oxidative damage, enabling corals to adapt to fluctuating light conditions and environmental stressors^2,3^.

During photosynthesis, the primary electron transport chain (ETC) is responsible for the efficient conversion of light energy into chemical energy. The main membrane-bound components of ETC are Photosystem II (PSII) and Photosystem I (PSI), as well as the cytb_6_f complex, which mediates the electron transport between the two photosystems together with the thylakoid-lumen-located plastocyanin (PC). Light-induced electron transfer in PSII drives water oxidation via the water oxidizing complex (WOC) with a tetra-manganese cluster as a catalytic site, which have various oxidation states, called S-states. The electrons liberated from water oxidation are transferred from the donor side of PSII, via the primary electron donor Chl assembly (P680), to the primary electron acceptor pheophytin (Phe), where from are transferred to a protein-bound plastoquinone molecule (Q_A_) acting as the first quinone electron acceptor^4^. Reduced Q_A_ can be reoxidized by a second plastoquinone electron acceptor called Q_B_, which is a mobile two-electron carrier. The double-reduced quinol form (Q_B_H_2_) can leave the binding site in PSII and deliver the electrons to a pool of PQ molecules located in the lipid phase of the thylakoid membrane^5^. From the PQ pool, the electrons are transferred via PC to the donor side of PSI, where they re-reduce the oxidized primary electron donor of PSI, called P700^+^, which is produced by light-induced charge separation in PSI, leading to the transfer of an electron to the acceptor side of PSI^6^. At the PSI acceptor side, ferredoxin (Fd) acts as a central hub for electron distribution towards downstream processes, primarily the production of NADPH via Fd-NAD(P)H-oxidoreductase (FNR)^7^. The electron transport through PSII and the cytb_6_f complex is coupled with trans-thylakoid proton translocation, and the produced proton gradient is utilized in ATP synthesis by the action of the membrane-bound ATP-synthase^8^. The NADPH and ATP molecules produced in the light-dependent phase of photosynthetic electron transport are utilized in various downstream processes, most importantly in CO_2_ reduction in the Calvin-Benson-Bassham (CBB) cycle, using CO_2_ as the final electron acceptor in the photosynthetic process^9,10^.

In addition to the above-described linear electron transport process, photosynthetic organisms have developed alternative electron transport pathways to adapt to varying environmental conditions and metabolic demands. These pathways include cyclic electron flow (CEF) around PSI, the alternative oxidase (AOX) pathway, the Mehler reaction, and chlororespiration. Each of these processes serves a distinct role in maintaining cellular energy balance, protecting against oxidative stress, and ensuring the flexibility and resilience of the photosynthetic apparatus. These alternative electron transport processes have been investigated in Symbiodiniaceae in earlier works^11–16^.

Kinetic measurements of variable Chl fluorescence, arising from PSII, provide a powerful tool to reveal the electron transport routes in Symbiodiniaceae and other microalgae. The yield of variable Chl fluorescence is regulated by the redox state of Q_A_, which is determined by the equilibrium of its reduction and reoxidation pathways. Importantly, Q_A_ can be reduced not only by forward electron transfer from the PSII reaction center, but also via charge equilibria with Q_B_ and the PQ pool^17^. Therefore, the Chl fluorescence yield is a sensitive indicator of the redox state of the PQ pool, which serves a central hub for different electron transport pathways. By using flash-induced chlorophyll fluorescence relaxation kinetics, a wave-like phenomenon was identified in Symbiodiniaceae under acute heat and microaerobic conditions, which is associated with the transient oxidation and re-reduction of plastoquinone (PQ) pool^18^. This Chl fluorescence wave was found to be associated with the operation of type II NAD(P)H dehydrogenase (NDH-2), indicating the existence of a slow, stromal-electron-source-related alternative electron transport^19^. The fluorescence wave components appearing in the several hundreds of milliseconds to a couple of seconds timeframe are informative about the alternative electron transport within the photosynthetic apparatus of algae cells^17,19–21^. The flash-induced fluorescence relaxation kinetics is a valuable indicator of the regulation of the electron transport network, and there is increasing evidence of its applicability in microalgae. The fluorescence wave phenomenon was also noted in intact coral species, such as *Pocillopora damicornis,* collected and measured at the Heron Island Research Station (unpublished data, Vass and Larkum, 2009), but the features and underlying mechanisms of the phenomenon were not investigated further. In the broader context of coral health, the flash-induced chlorophyll fluorescence patterns could provide insight into the complex interplay of the electron transport pathways between the coral host and the symbiotic algae. Since the fluorescence relaxation can be measured in extended timescales (several hundreds of seconds), it is imperative to study the changes in fluorescence relaxation that are related to the slow electron transport processes leading to the reduction of the PQ pool.

In addition to alternative electron transport routes that take place within the cells, some unicellular photosynthetic (and non-photosynthetic) organisms are able to deliver electrons to acceptors located outside the cells in a process denoted extracellular electron transport (EET)^22,23^. In photosynthetic microorganisms such as cyanobacteria, high light intensity or low carbon availability leads to over-reduction of the primary electron transport chain of the thylakoid membrane, resulting in photodamage^24,25^. One potential protective mechanism to counteract this effect is utilizing EET to offload surplus electrons to external acceptors, thereby protecting the photosynthetic machinery. Electrons from the PQ pool (PQH_2_) can be transferred to cytoplasmic carriers, including ferredoxins and soluble cytochromes, or directly to extracellular components via some unidentified proteins^26^. The involvement of conductive pili was also suggested in this process in cyanobacteria^27^. In some species, membrane-bound cytochromes and other transporters, such as riboflavin in the plasma membrane, can mediate electron transfer to extracellular components^28,29^. This mechanism not only protects the photosynthetic machinery but also allows these microorganisms to engage in ecological interactions, such as reducing external compounds, e.g., Fe^3+^, or facilitating electron exchange with other microbes in biofilms or mats^30^. By integrating EET into their metabolic networks, photosynthetic microorganisms enhance their adaptability to dynamic environmental conditions, sustaining energy production while minimizing photodamage. Interestingly, some microbes are able not only to deliver electrons from the cells to extracellular acceptors but also to extracellular electron uptake (EEU) from sources outside the cells^31,32^.

A key aspect of coral symbiosis is the finely tuned exchange of metabolites between the coral host and its algal symbionts. Once inside the host’s gastrodermal cells, the algae are enclosed within symbiosomes, specialized intracellular compartments that create a controlled microenvironment. The symbiosome maintains an acidic pH (∼4) through active proton pumping, which enhances carbon dioxide availability for photosynthesis^33,34^. The transfer of essential compounds between the host and Symbiodiniaceae requires transport across multiple membranes, including the algal plasma membrane, the symbiosome membrane, and the host cell membrane. Several transporters facilitate the movement of critical nutrients, such as nitrogen, carbon, and phosphorus, as well as cofactors like riboflavin and folate, which are essential for cellular metabolism and detoxification^29^. Notably, riboflavin transporters have been linked to EET in bacteria^28^, and their presence in the plasma membrane of *Symbiodinium*^29^ raises the intriguing possibility that similar electron transfer mechanisms may exist in coral-algal symbiosis. While EET in coral systems remains largely unexplored, its potential role in stress adaptation and metabolic regulation demands further investigation.

In coral reef ecosystems, where light intensity and temperature fluctuations influence photosynthetic activity, EET may serve as a crucial adaptation for maintaining redox homeostasis. Symbiotic microorganisms associated with corals, including photosynthetic algae and bacteria, could use EET as a safety valve to export excess electrons to extracellular acceptors when primary electron sinks, such as NADP⁺ or carbon fixation pathways, become overwhelmed. This process could help in reducing oxidative stress, which is a major factor contributing to coral bleaching under environmental stress. Recently, Marcel et al. (2025) demonstrated using a bio-electrochemical system that *Symbiodinium* cells in culture exhibited an efficient EET, about 2.5-times higher than diatoms^35^. As climate change and anthropogenic pressures continue to threaten coral ecosystems, investigating the role of EET in these systems could provide new insights into their adaptive strategies. Understanding how microbial EET contributes to coral-algal partnerships and broader reef ecosystem functions may be essential for developing conservation approaches to sustain reef health in changing ocean conditions.

The aims of the current work were i) to study the wave phenomenon of flash-induced Chl fluorescence relaxation in intact corals in the long 10-400 second timeframe, and to reveal its link to the electron transport pathways and the overall stress response in the coral host and the algae, and ii) to clarify the existence of EET in the coral-Symbiodiniaceae system. Our data show that corals display a slow-wave phenomenon in flash-induced Chl fluorescence relaxation that is associated with the redox level changes of Q_A_ and influenced by the ambient oxygen level via respiration of the coral host. The fluorescence characteristics together with the reduction of the extracellular electron acceptor FeCN suggest that electron transfer occurs from the interior of symbiont cells to the extracellular space, and possibly also in the reverse direction.

## Results

### Flash-induced chlorophyll fluorescence relaxation kinetics in the minutes time scale

Application of a single-flash excitation results in a jump of the variable Chl fluorescence yield in cultured *Fugacium* cells under enzymatically induced microaerobic conditions, due to the light-induced reduction of Q_A_, which is followed by a biphasic decay on the hundreds of μs to tens of ms timescale, reflecting reoxidation of Q_A_^−^ as observed in other microalgae^17,21^. Interestingly, this decay is followed by a gradual, slow rise of fluorescence yield, forming a wave-like pattern over a period of 10-400 seconds (Fig. 1A). The phenomenon is similar to that observed earlier in Symbiodiniaceae under heat and microaerobic conditions^18^. However, the kinetics are quite different since the rapid rise in the fluorescence yield took place already from a few milliseconds and declined back to the baseline after 100 s when heat and microaerobic conditions were combined^18^, in contrast to the slow fluorescence rise observed here. Importantly, this pattern of slow fluorescence rise was also found in intact coral nubbins (*Pocillopora*) as shown in Fig. 1B. However, in this case, the fluorescence yield after 400 s was significantly higher than the initial baseline, while in the *Fugacium* cells the 400 s fluorescence yield reached practically the initial baseline level (Fig. 1A). The application of DCMU, an inhibitor of electron transfer between the Q_A_ and Q_B_ electron acceptors in PSII, resulted in the complete elimination of the fast and intermediate phases of fluorescence relaxation, with only the slow phase remaining, which originates from the charge recombination of Q_A_^-^ and the S_2_ state of the water-oxidizing complex^17^. Furthermore, the DCMU treatment also led to the disappearance of the fluorescence wave, both in the cultured cells and intact coral nubbins (Fig. 1A and B), which suggests that charge equilibrium between Q_A_^-^ and Q_B_, and the PQ pool is a crucial factor in the formation of this phenomenon.

**Figure 1:**
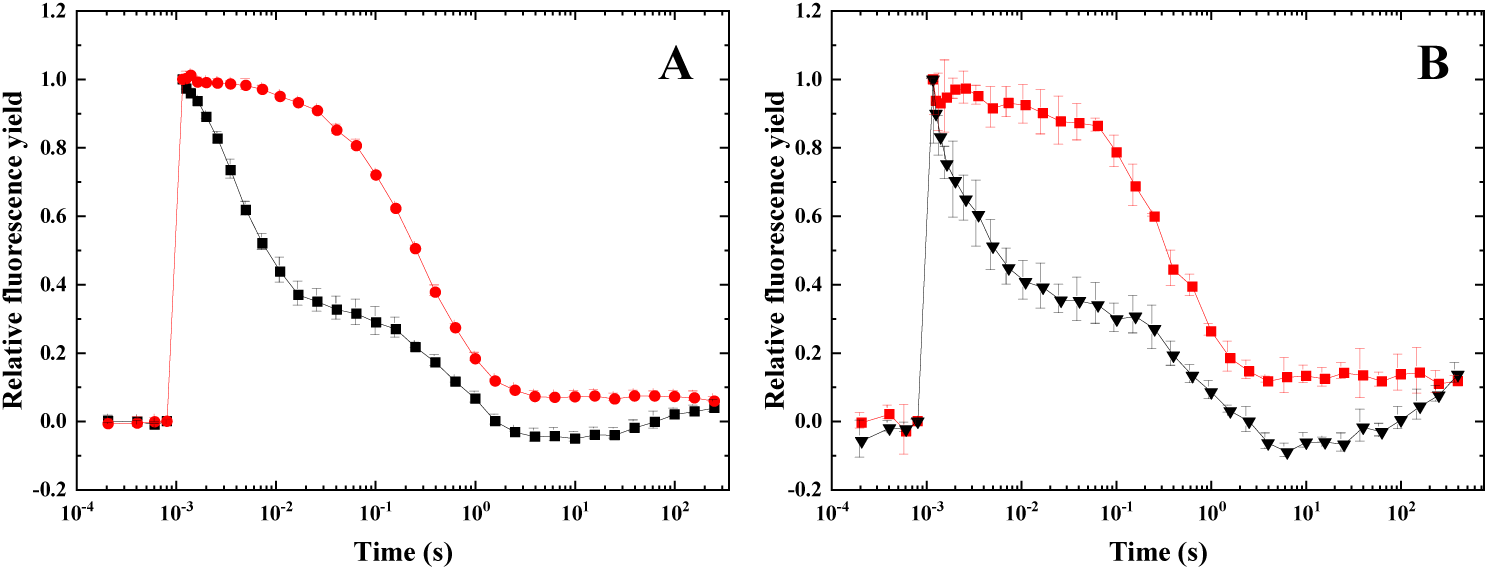
Flash-induced Chl fluorescence yield changes. (A) *Fugacium kawagutii* (CS156), (B) *Pocillopora damicornis* were measured under enzymatically induced microaerobic conditions, in the absence (black) and presence of 20 μM DCMU (red). The curves are mean values from 3 biological replicates and shown after normalization to the same initial amplitude.

A similar slow rise in Chl fluorescence over a period of 10-400 seconds was also observed in other coral species, including *Acropora* and *Seriatopora*. This rise in fluorescence was also blocked in the presence of DCMU, as observed in *Pocillopora*, thus indicating its uniformity across species (Fig. 2).

**Figure 2:**
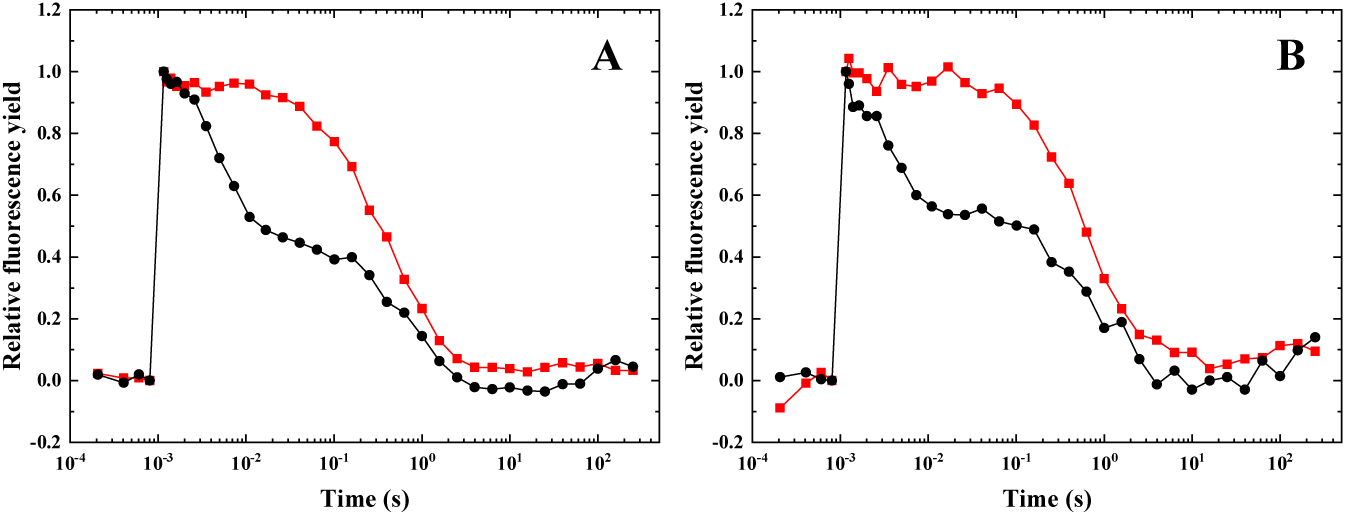
Flash-induced fluorescence yield changes in corals. (A) *Acropora millepora*, (B) *Seriatopora hystrix* were measured under enzymatically induced microaerobic conditions, in the absence (black) and presence of 20 μM DCMU (red). The curves are shown after normalization to the same initial amplitude.

To determine if the observed slow fluorescence wave is induced by the actinic flash, and if the wave phenomenon is associated with the redox reactions of the formed Q_A_, measurements were conducted with and without the application of the actinic flash. In the absence of the actinic flash, the baseline fluorescence level was practically constant up to 400 s in the *Fugacium* cells (Fig. 3A). In contrast, in the coral nubbins, a small, but clearly observable increase of baseline fluorescence was observed, which became obvious after approximately 100 seconds in the logarithmic timescale (Fig. 3B). This observation shows that in the intact corals Q_A_ is gradually reduced in the dark, which effect was absent in cultured *Fugacium* cells. Dark reduction of Q_A_ in the host-embedded symbionts can occur due to electron transfer to Q_A_, via charge equilibrium with the PQ pool, which in turn can be reduced via an interaction of the symbiont and the coral host. This interaction may arise from a metabolic effect, such as oxygen consumption by the host, or from an electron transfer between the host and the symbiont. It is also of note that the fluorescence dip was not observed in the absence of actinic flash and it appeared only in the flash-illuminated samples (Fig. 3).

**Figure 3:**
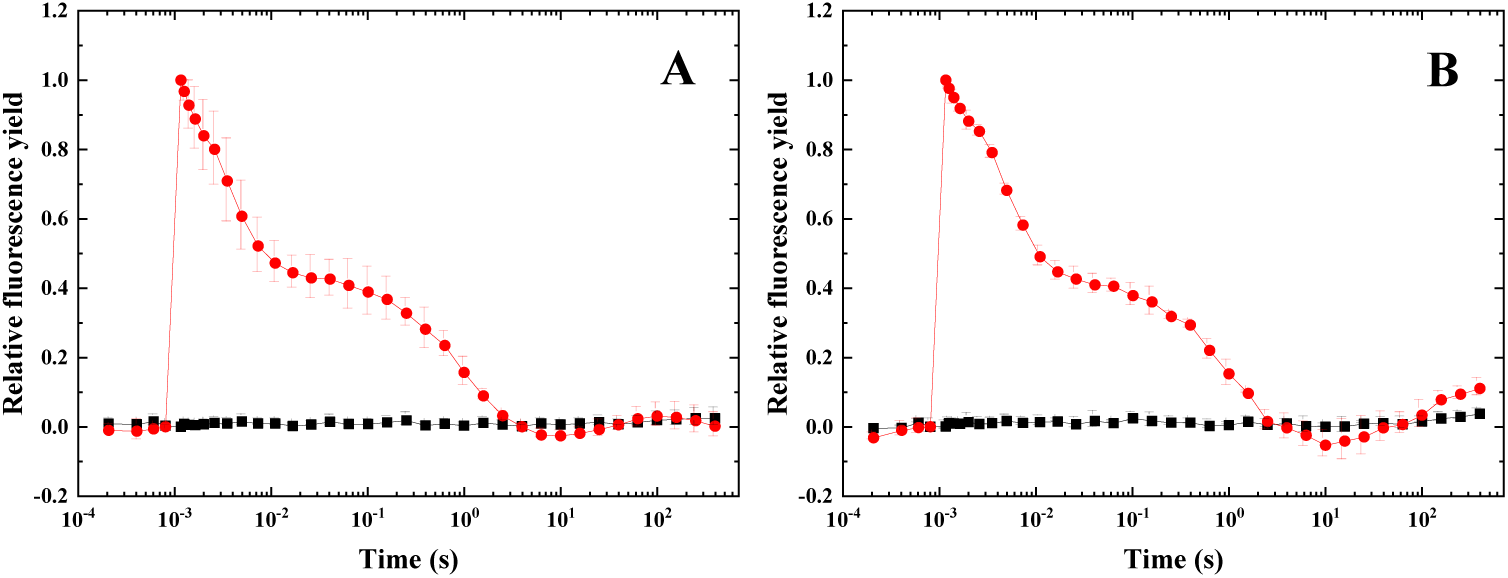
Fluorescence yield changes in the dark and after flash excitation. (A) *Fugacium kawagutii* (CS156), (B) *Pocillopora damicornis* were measured under enzymatically induced microaerobic condition, in the absence (black) and presence of actinic flash (red). The curves are mean values from 3 biological replicates and shown after normalization to the same initial amplitude.

### Oxygen-dependent change in fluorescence relaxation kinetics of corals

The fluorescence wave in the time range of 10-400 seconds was observed earlier in only under microaerobic conditions in *Fugacium*^18,21^, indicating that oxygen-limitation-induced reduction of the PQ pool was required for the slow fluorescence rise^18,21^. However, in coral nubbins, kept in artificial seawater in the relatively small volume (2 mL) of the measuring cuvette, the oxygen level can spontaneously decrease due to the respiration activity of the coral host, which can also consume oxygen from the surrounding liquid medium. During 400 s incubation of the coral nubbin in the cuvette, the dissolved O_2_ level dropped from 230 μM/L to 135 μM/L, while during the continued incubation of the same coral nubbin, the O_2_ level dropped further to 50 μM/L (Fig. 4A, black and red curves, respectively). Under these conditions, the baseline fluorescence level showed a gradual increase in the absence of actinic flash in the dark (Fig. 4B, black curve). During the repeated measurement, the baseline fluorescence increased further from the level reached at the end of the first period (Fig. 4B, red curves), demonstrating the gradual reduction of Q_A_ in the host-embedded symbiont. When the coral samples were made microaerobic by an oxygen-consuming enzyme treatment, the level of dissolved oxygen dropped to practically zero (ca. 1 μM/L) and did not change further during the 400 s incubation period (Fig. 4A, blue curve). Interestingly, the baseline fluorescence showed a continued increase during the microaerobic incubation (blue curves in Fig. 4B), even though the level of dissolved oxygen did not decrease further (blue curve in Fig. 4A). However, the extent of F_o_ rise was smaller under enzymatic O_2_ removal than under spontaneous O_2_ consumption by the coral host (blue vs. red and black curves in Fig. 4B) than occurred under respiratory oxygen uptake by the coral host (Fig. 4B). When the coral nubbins, which were incubated in the cuvette without any addition, were illuminated with an actinic flash, a gradual slowdown of the fast phase of the Chl relaxation was observed with a concomitant increase of the middle phase, demonstrating the gradual reduction of the PQ pool (Fig. 4D, E). Enzymatic removal of oxygen resulted in further slowdown of the fast phase and an increase of the middle phase, showing a more complete reduction of the PQ pool and a higher level of Q_A_ reduction. After prolonged incubation of the nubbins in the cuvette without any addition, a clear fluorescence wave phenomenon was observed (Figs. 4E), even though only the respiration of the host consumed the dissolved oxygen without artificial oxygen removal treatment. Importantly, the wave became more pronounced when the dissolved oxygen level dropped close to 0 after enzymatic oxygen removal (Fig. 4F). These observations demonstrate that spontaneous oxygen consumption by the host respiration can induce the fluorescence wave, which therefore provides a useful indicator of the redox state of the electron transport chain of the symbiont *in hospite*.

**Figure 4:**
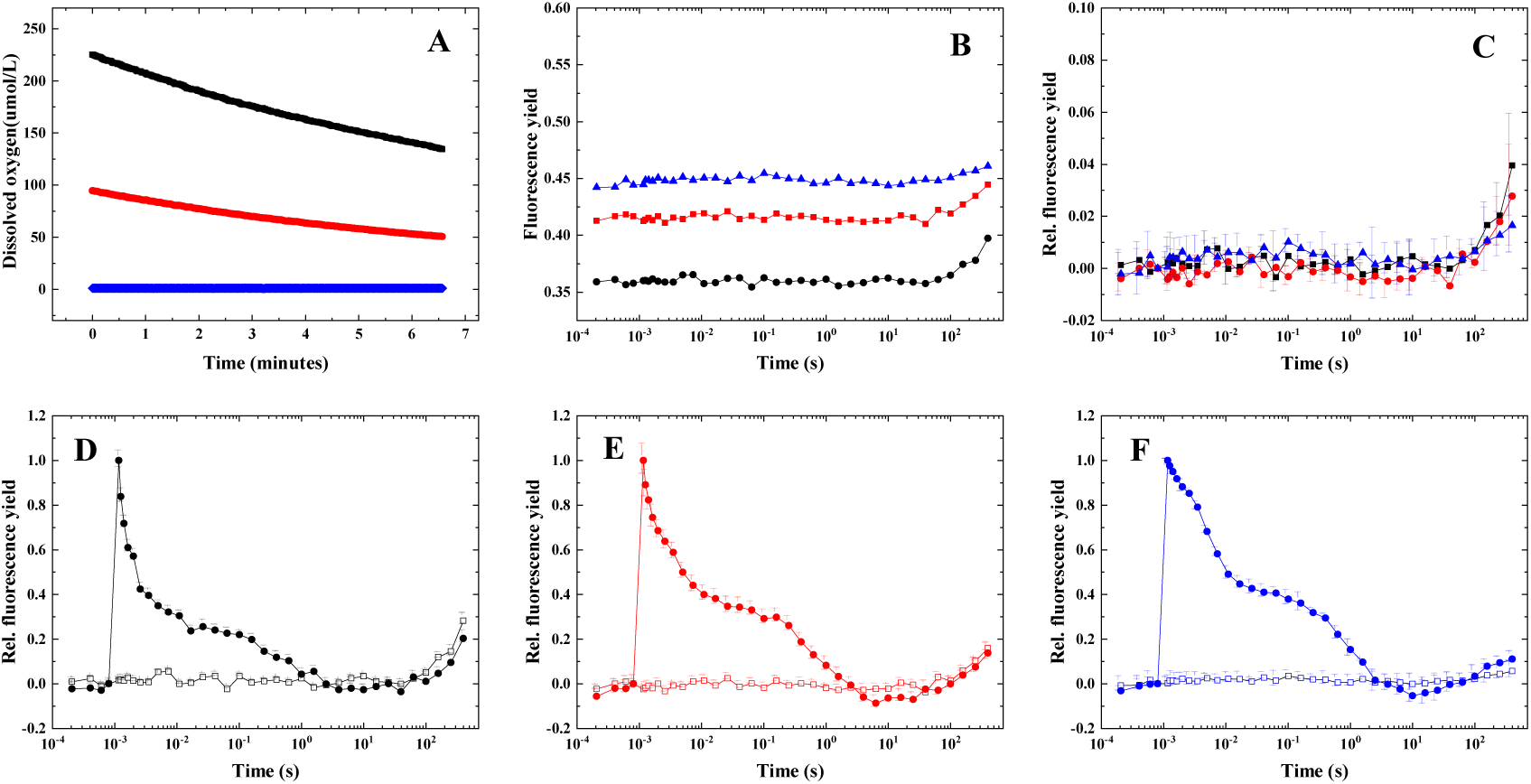
Effect of dissolved oxygen levels on Chl fluorescence kinetics of Pocillopora damicornis. Coral nubbins were placed into a 1 cm cuvette and were incubated in artificial seawater without addition, or in the presence of oxygen-consuming enzyme mix. Changes in dissolved oxygen level were followed by an optical oxygen sensor. Chl fluorescence measurements were done during incubation under the respective conditions. (A) Dissolved oxygen level changes induced by host respiration (black, red), or by enzymatic oxygen removal (blue) in darkness. (B) F_o_ Chl fluorescence changes without actinic flash under the conditions shown in panel A. (C) F_o_ fluorescence changes shown after normalization to identical initial values. Panels D, E, and F show Chl fluorescence yield changes after single turnover flash excitation (closed symbols) and without flash (open symbols) under the oxygen consumption conditions shown in Panel A, i.e. induced by host respiration (black and red) and enzymatic oxygen removal (blue). The curves in Panels D-F are normalized to the same initial amplitudes. The curves are mean values from 3 biological replicates.

It is worth noting that the initial observation of the slow-wave Chl fluorescence phenomenon in corals was made during a research trip to the Heron Island Research Station in 2009. When freshly collected coral samples were placed in the measuring cuvette, where host respiration could consume most of the dissolved oxygen, essentially the same Chl fluorescence wave characteristics were observed as shown here (Vass and Larkum, unpublished, but see Fig. 5).

**Figure 5.**
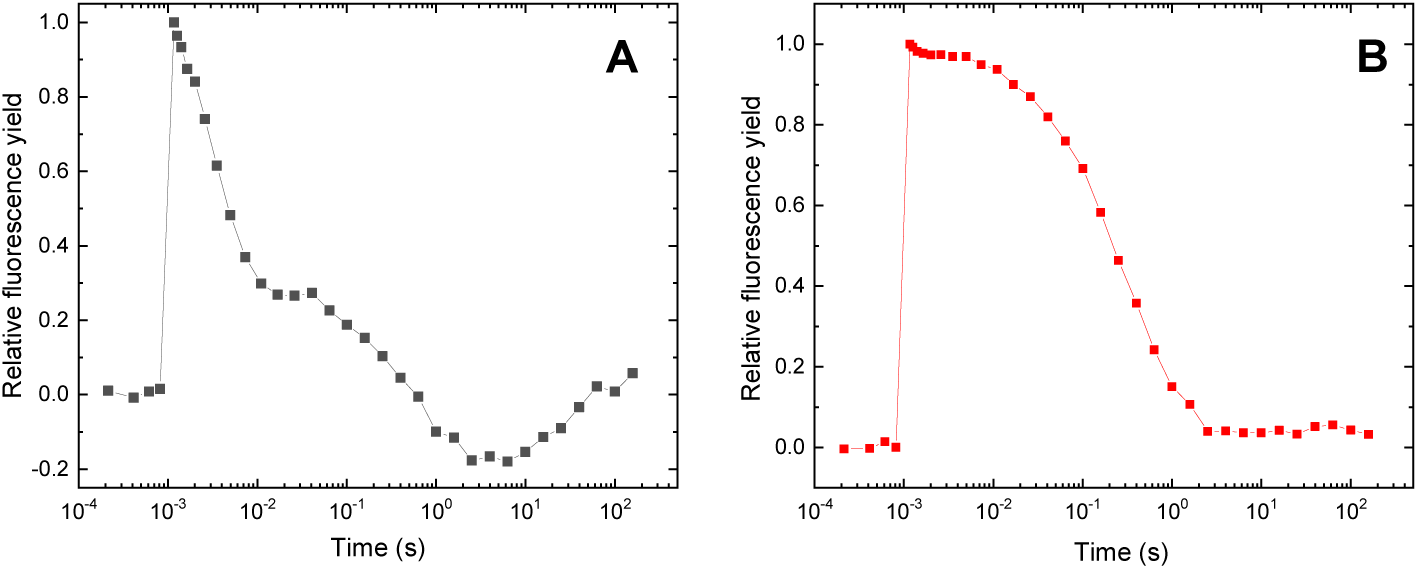
Flash-induced fluorescence relaxation in *Pocillopora damicornis* in the absence (A) and presence (B) of 10 μM DCMU. Coral samples were collected at Heron Island lagoon on the Great Barrier Reef of Australia (151°550E, 23°270S) in November 2009. The colonies were maintained in circulating natural seawater at the Heron Island Research Station in 500 L aquaria under natural sunlight, attenuated by a shade cloth. Flash-induced Chl fluorescence was measured in coral nubbins placed in a 1 cm glass cuvette, as described in the Materials and Methods, using the same type of PSI instrument as in the present study, except a shorter time window: 214 μs-158 s, instead of 150 μs-400 s used here.

### Effect of ferricyanide on Chl fluorescence kinetics

The characteristics of the above-described slow Chl fluorescence wave in intact coral nubbins strongly indicate that this phenomenon is associated with electron transport interaction between the algal symbiont and the coral host. To get further insight into this interesting effect, an artificial acceptor, 1 mM potassium ferricyanide (FeCN), which does not penetrate the cell wall of the symbiont, was employed to elucidate whether extracellular electron transport modulates the slow wave phenomenon in Symbiodiniaceae. FeCN was observed to not only eliminate the slow wave (10–400 s) of fluorescence relaxation but also to reduce the middle phase of the fluorescence decay (5–15 ms) in *Fugacium* (Fig. 6A), as well as the initial F_o_ fluorescence yield (Fig. 6A inset). These effects strongly indicate that the extracellular electron acceptor, FeCN, can receive electrons from the interior of the *Fugacium* cells, which directly or indirectly originate from the PQ pool, demonstrating the existence of EET in *Fugacium*.

**Figure 6:**
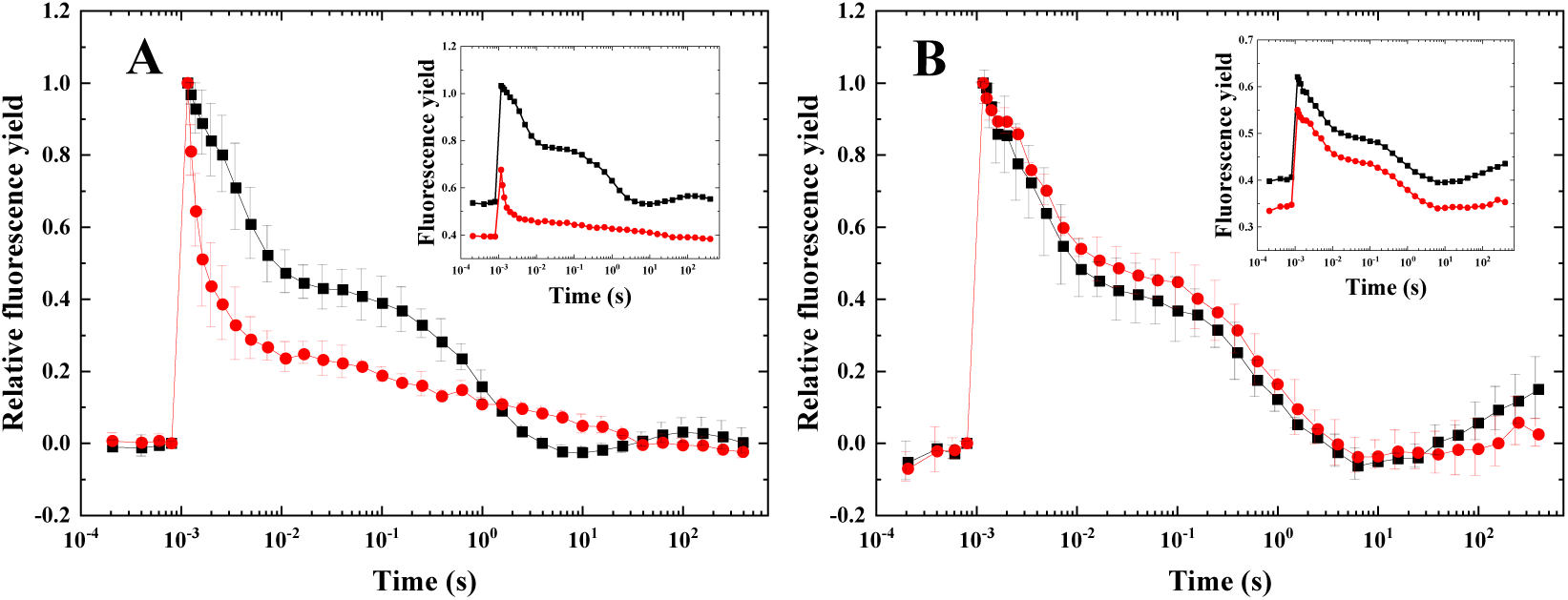
Effect of FeCN on flash-induced fluorescence relaxation. (A) *Fugacium kawagutii* (CS156), (B) *Pocillopora damicornis* under enzymatically induced microaerobic conditions, in the absence (black) and presence of 1 mM FeCN (red). The curves are normalized to the same initial amplitudes, but the insets show the curves without normalization. The curves are mean values from 3 biological replicates.

In intact coral nubbins, FeCN significantly decreased F_o_ and slightly reduced the fluorescence rise over extended periods (10–400 s), as shown in Fig. 6B, suggesting a FeCN-induced compensation of host-induced reduction of Q_A_ in the symbiont. However, FeCN did not affect the middle phase (5–15 ms) of fluorescence relaxation. The significant decrease of the F_o_ level in the absence of actinic flash (Fig. 7A) indicates that, when photosynthetic electron transport is inactive, electrons are drained from the PQ pool to FeCN. However, the remaining F_o_ rise in the dark (Fig. 7A, B) shows that a large part of the host-induced reduction of Q_A_ exists even in the presence of FeCN. It is also of note that DCMU, which inhibits electron transfer between Q_A_ and Q_B_, completely blocked the F_o_ rise in the dark, which demonstrates that this effect reflects dark reduction of Q_A_ from the PQ pool via Q_B_.

**Figure 7:**
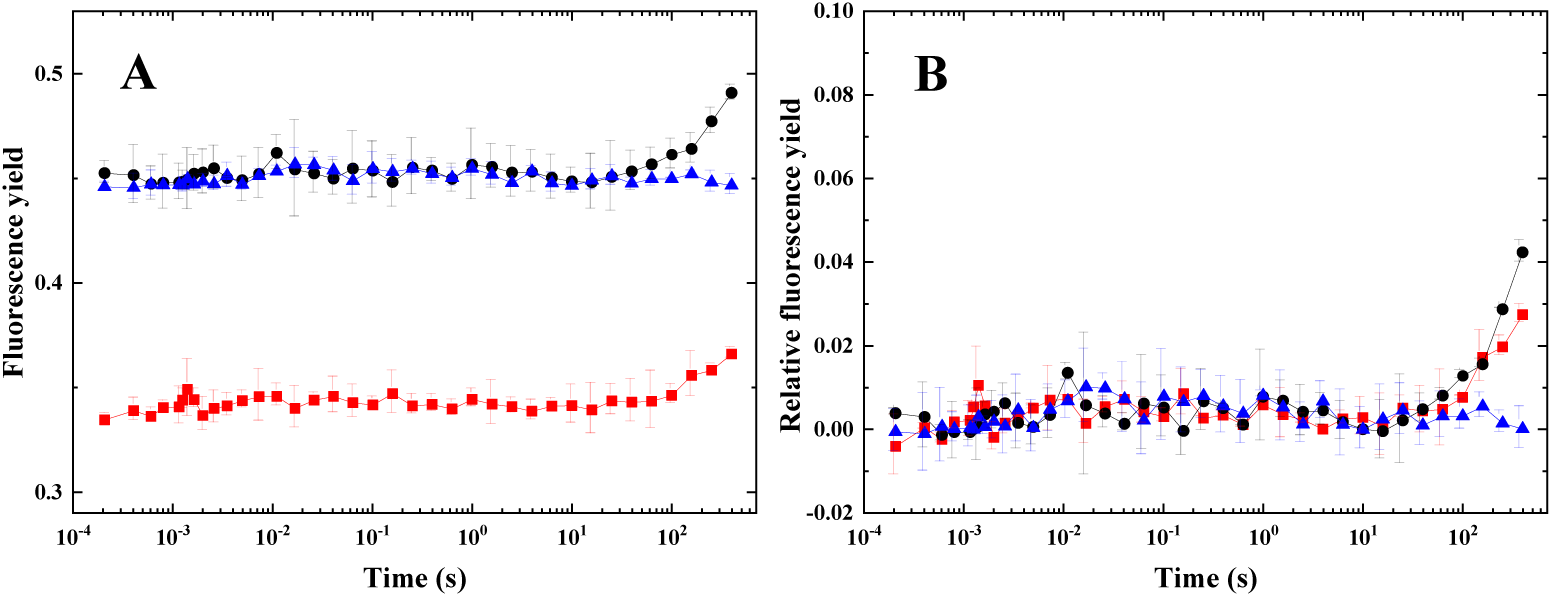
Effect of FeCN on the F_o_ fluorescence yield in the dark. Fluorescence yield changes were measured without actinic flash in *Pocillopora damicornis* under enzymatically induced microaerobic conditions. (A) raw data, (B) after normalization to the same initial fluorescence levels. Samples were kept in the dark in the absence (black) and presence of 1 mM FeCN (red), the blue curve shows the effect of 10 μM DCMU in the absence of FeCN. The curves are mean values from 3 biological replicates.

Variable Chl fluorescence measurements under continuous illumination (OJIP transients) provide information on electron transport in PSII, the PQ pool, and beyond. In *Fugacium kawagutii* cultures, FeCN caused only a slight retardation of the fluorescence rise from the J phase (Fig. 8A and B), indicating that FeCN-dependent electron extraction from the cells does not compete efficiently with light-induced reduction of the electron transport chain on the timescale of the fluorescence transient measurements. The rise of the variable fluorescence in the coral nubbins was significantly slower than in the isolated *Fugacium* cultures, but reached the same maximal P value. This delayed fluorescence rise in the coral nubbins shows a less efficient reduction of the electron transport carriers than in the isolated symbiont cells, which may indicate the presence of an alternative electron transfer process that competes with light-induced reduction of the electron transport chain. In the presence of FeCN, the fluorescence rise kinetics were further retarded in the coral nubbins from the J phase (Fig. 8B), indicating a FeCN-dependent electron transport to the exterior of the cells. Although the cell wall of the symbionts is more penetrable *in hospite* than in the intact cells, the fluorescence rise kinetics are clearly different from that observed in isolated thylakoids (Fig. 8B inset), where FeCN can directly accept electrons from PSII, and retards fluorescence rise even before the J phase. This comparison shows that FeCN cannot interact directly with PSII and the PQ pool in the host-embedded symbiont cells, and can accept electrons only from the exterior of the symbiont cells.

**Figure 8:**
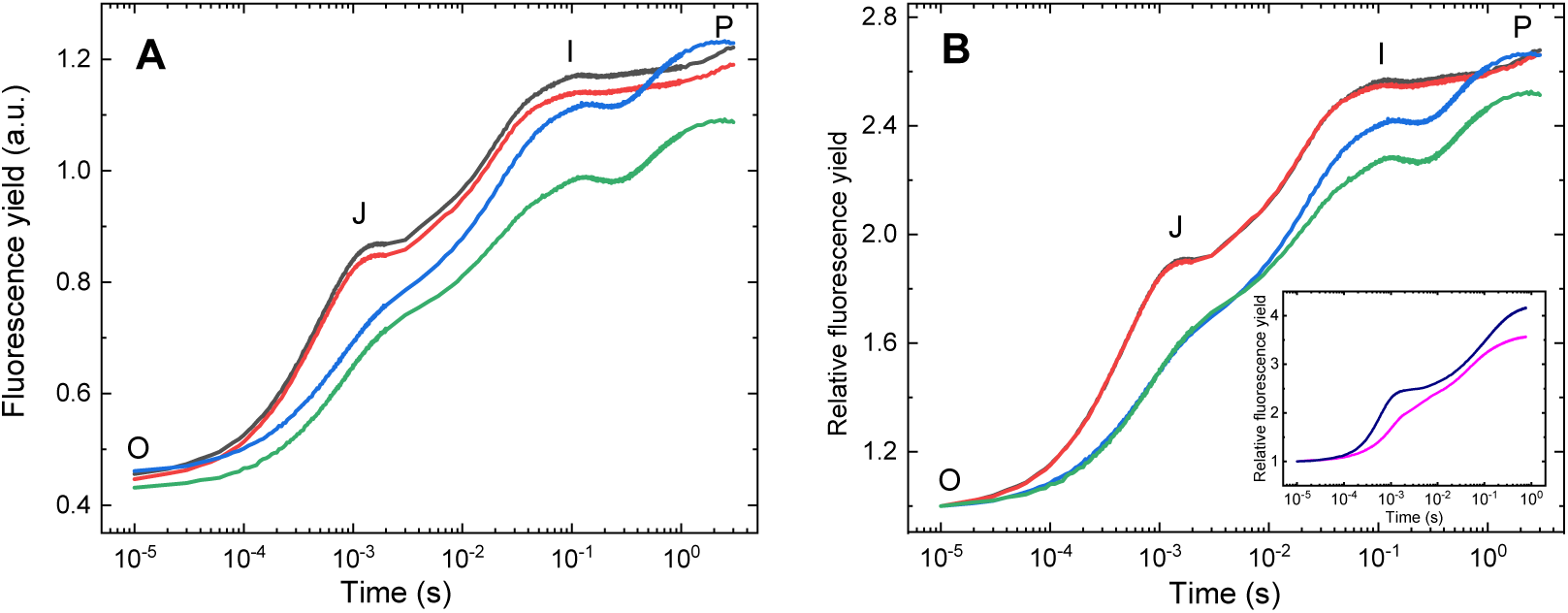
Effect of FeCN on the OJIP-type Chl fluorescence yield traces of *Fugacium kawagutii* and *Pocillopora damicornis*. Variable Chl fluorescence was measured under continuous illumination in *Fugacium kawagutii* (CS156) cultures (black and red curves), *Pocillopora* nubbins (blue and green curves), and isolated spinach thylakoids (Inset, navy and magenta curves). The samples were measured without addition (black, blue, and navy) or in the presence of 1 mM FeCN (red, green, and magenta). The curves are mean values from 3 biological replicates, and are shown without normalization (A) and after normalization to identical initial, O, values (B and inset).

### Ferricyanide reduction assay

To verify the existence of electron transport directed outwards of *Fugacium* cells, as indicated by the above fluorescence data, a FeCN reduction assay was conducted, which is based on the measurement of absorbance at 420 nm reflecting the reduction of ferricyanide to ferrocyanide^23^. Since FeCN molecules are soluble artificial redox mediators, impermeable to the plasma membrane^23^, the observation of FeCN reduction shows that electrons from the interior of cells can reach the extracellular environment. The assay was performed under a variety of conditions, including those involving isolated *Fugacium* cultures, *Fugacium* cells whose cell wall was partially digested (mimicking the symbiotic lifestyle), and intact coral nubbins. The results demonstrated a notable reduction of FeCN in both intact cells and cells with partially digested cell walls when the samples were kept in growth light (50 µmol photons m^-2^ s^-1^) for two hours. DCMU had only a small inhibitory effect on the reduction process (not shown in the figure), suggesting that most of the electrons that reduce FeCN are derived from light-independent processes. In agreement with this, the dark reduction process of FeCN continued for ca. 15 hours until the whole amount of added 1 mM FeCN was completely reduced (not shown).

Reduction of FeCN was also observed in the presence of intact coral nubbins (Fig. 10A). To account for variations among the different coral samples, the reduction rates were normalized relative to the weight of the coral nubbins, showing comparable results across samples (Fig. 10B).

To compare the FeCN reduction rates among the different sample types, we normalized the rates to equal Chl amounts by assuming that Symbiodiniaceae cells have the same oxygen evolving capacity in isolated cultures and *in hospite*, and therefore the measured oxygen evolution rates are proportional to the total Chl content in both the isolated call cultures and the coral nubbins. Based on this assumption, the measured O_2_ evolution rates of the coral nubbins were converted into estimated total Chl amounts in the same nubbins. A comparison of FeCN reduction rates with respect to Chl concentrations revealed that intact *Fugacium* (CS156) cells exhibited approximately 13.5 ± 2.4 µmol mg Chl^-1^ h^-1^; however, the rate showed variation among different culture batches. The rate was somewhat higher in CS156 cells with partially digested cell wall, at 15.8 ± 5.3 µmol mg Chl^-1^ h^-1^. The estimated FeCN reduction rate of intact *P. damicornis* coral nubbins was approximately 29.5 ± 7 µmol mg Chl^-1^ h^-1^. The apparently elevated rates observed in intact corals as compared to those in *Fugacium* cells in culture may be attributed to the distinct metabolic conditions of the symbiont cells within the coral tissue, especially the availability of CO_2_ as a final electron acceptor for O_2_ evolution that could alter the estimation of Chl amount in the coral nubbins based on the O_2_-evolving capacity. However, some direct contribution of the host to the extracellular electron transfer processes cannot be excluded either.

## Discussion

### Flash-induced chlorophyll fluorescence relaxation as a stress marker in corals

Flash-induced Chl fluorescence relaxation kinetics provides useful information about a range of photosynthetic electron transport processes, including linear and alternative electron transport pathways and charge recombination^36^. The redox state of Q_A_ regulates Chl fluorescence yield: In the presence of reduced Q_A,_ the primary charge separation between P680* and Phe is suppressed, therefore, the excitation energy of P680* is dissipated in the form of fluorescence, resulting in high Chl fluorescence yield^37^ (Scheme I).

Earlier, we investigated the photosynthetic electron transport processes in Symbiodiniaceae, utilizing the features of the Chl fluorescence wave phenomenon, which appears in the time range from tens of milliseconds to tens of seconds, and was found to be associated with the NDH-2-mediated alternative electron transport pathway^19^. In the current work, we observed that under microaerobic conditions, the flash-induced Chl fluorescence relaxation exhibits characteristic features, which are like the previously described wave phenomenon, but occur in a much longer (10-400 s) time scale in both cultured *Fugacium* cells and intact coral nubbins across multiple coral species, such as *Pocillopora*, *Acropora,* and *Seriatopora*. The early observations (Vass and Larkum, 2009, Fig. 5) together with the present data demonstrate that the slow-wave fluorescence phenomenon is an intrinsic characteristic of fully active coral specimens when they are grown either in the laboratory or under natural conditions in the ocean, and facilitate a deeper understanding of photosynthetic and other electron transport processes within the Symbiodiniaceae, and between the Symbiodiniaceae and their coral host.

The basic difference between the slow fluorescence wave observed here and the earlier reported fast wave in cultured Symbiodiniaceae cells is that the dip phase, i.e. the decline of the fluorescence intensity below the initial baseline, appears ca. 10 s after the actinic flash in the slow wave (Figs. 1 - 4), in contrast to 50-100 ms observed in the fast wave^19^. Similarly, the bump, i.e. the maximal level of the fluorescence rise, is observed after 100-200 s in the slow wave, in contrast to 10-20 s in the fast wave^19^. The formation of the fast fluorescence wave in cultured Symbiodiniaceae was explained by the transient oxidation of the PQ pool driven by enhanced PSI electron uptake, which becomes more prominent when PSII activity is inhibited by heat stress. Heat also suppresses Calvin–Benson–Bassham (CBB) cycle activity, leading to stromal over-reduction and altered electron partitioning, potentially increasing electron flow into the PQ pool through alternative pathways, such as via NDH-2^19^. Under microaerobic conditions, reduced availability of O₂ limits PTOX activity, causing the PQ pool to remain more reduced. This slows down Q_A_⁻ reoxidation and also facilitates the backflow of electrons from the PQ pool to Q_A_^-^, which in turn modulates the Chl fluorescence yield (see Scheme I).

The elimination of the slow fluorescence wave in *Fugacium* cultures and in the various coral species (Figs. 1 and 2) by DCMU, which blocks electron transport between the Q_A_ and Q_B_ electron acceptors in PSII (Scheme I), indicates that the slow wave is related to the operation of the linear electron flow. Furthermore, the observation that the dip appears only when an actinic flash is applied—and is absent without actinic illumination (Fig. 3)—supports the conclusion that this feature arises from photosynthetically driven electron-transport processes. The significantly delayed appearance of the dip in the slow-wave conditions can be explained by the presence of fully active PSII, which can compensate for a longer time the PQ pool oxidation effect by PSI, as occurs in heat-treated samples^19^. Similarly, it takes a longer time for the reverse electron flow from the stromal components to induce a reduction effect on the PQ pool when the CBB cycle is fully active, in contrast to heat-induced inhibition of the CBB cycle^19^.

**Scheme I.**
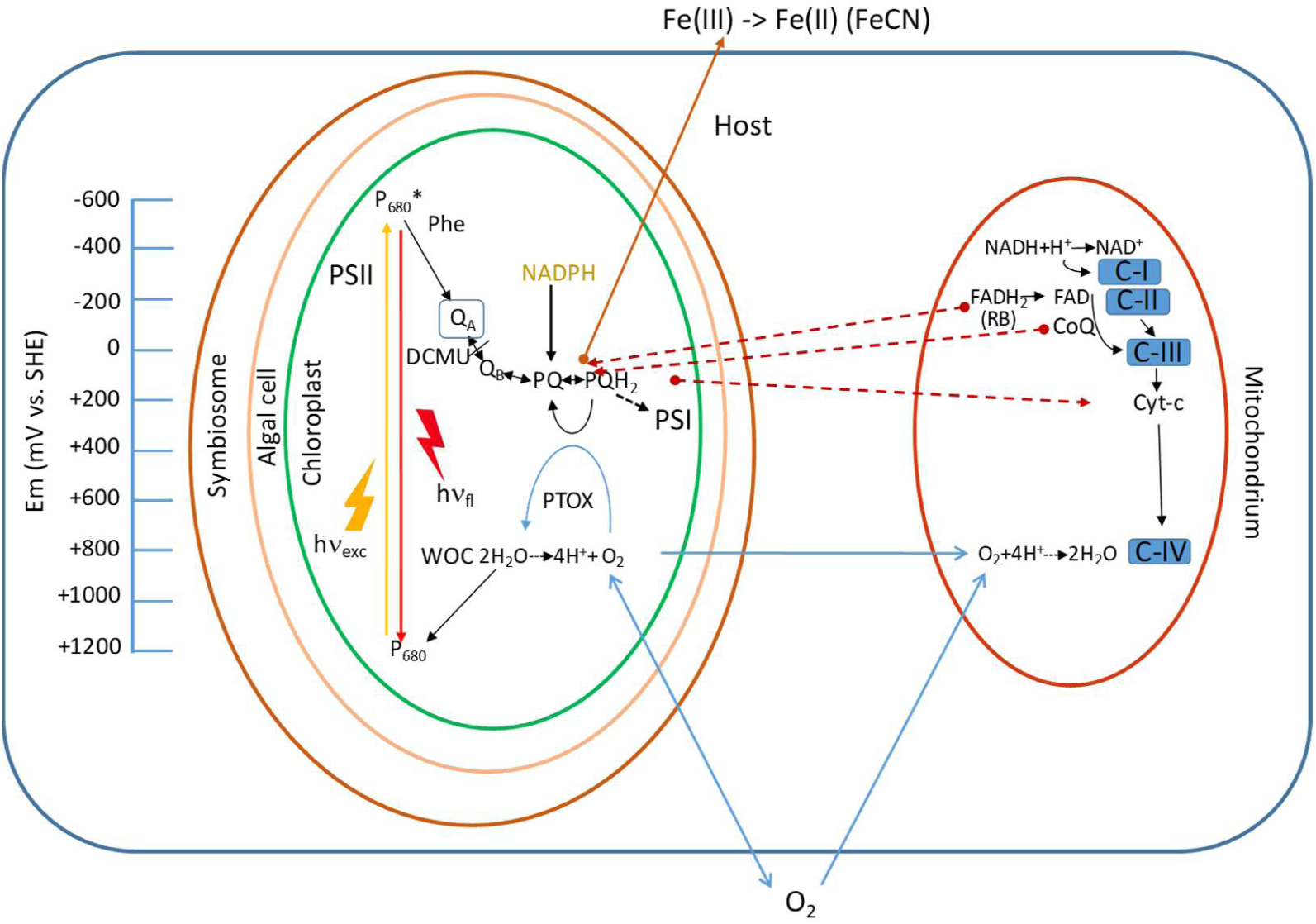
Interaction of photosynthetic electron transport in the symbiont with the respiratory electron transport in the host, in coral symbiosis. The figure shows the initial components of light-induced electron transport in the Photosystem II (PSII) complex and the PQ pool located in the thylakoid membranes of the symbiont’s chloroplasts, as well as the main components of the respiratory electron transport chain located in the mitochondria of the host cells. The yellow and red arrows show light-induced excitation of the PSII reaction centre Chl P680, and the fluorescence emission from the excited Chl, respectively. The black arrows show electron transport processes, the blue arrows represent O_2_ equilibria among the chloroplast, host mitochondria and the environment, the solid brown arrow shows extracellular electron transfer from the chloroplasts to extracellular electron acceptors, such as FeCN and Fe(III), while the dashed brown arrows indicate the putative extracellular electron transfer processes between the chloroplast of the symbiont and the mitochondria of the host. Light absorption induces the excitation of the special Chl assembly, P680, in PSII, which initiates charge separation between P680 and Phe, followed by electron transfer from Phe to the Q_A_ and Q_B_ quinone electron acceptors leading to the accumulation of reduced PQ (PQH_2_) in the so-called PQ pool in the lipid phase of the thylakoid membrane. (Electrons from the PQ pool are transferred to Photosystem I via the cytb_6_f complex and plastocyanin, resulting in NADPH production, not shown in the figure.) PQ can also be reduced to PQH_2_ via chlororespiration. During this light-independent process, electrons are transferred from NADPH to PQ, via the NDH-2 complex located in the thylakoid membrane. PQH_2_ can be oxidized back to PQ via the so-called chloroplast terminal oxidase (PTOX), transferring the electron to O_2_ to form H_2_O. Thermally activated charge equilibria from the PQ (PQH_2_) pool via Q_B_ can re-populate Q_A_^-^, which is the primary regulator of the yield of variable Chl fluorescence, since in the presence of Q_A_^-^ the primary charge separation is suppressed, resulting in the dissipation of the excitation energy of P680 in the form of fluorescence emission. Therefore, the reduction level of the PQ pool can be monitored by measuring the variable Chl fluorescence yield. The main components of the respiratory electron transport in the mitochondria of the host cells are the so-called Complex I,.., Complex IV (abbreviated with C-I,.., C-IV). C-I oxidizes NADH, which is supplied from the cellular metabolism via the citric acid cycle, to NAD^+^ and reduces ubiquinone (CoQ). C-II transfers electrons from succinate dehydrogenase via FADH_2_ to CoQ. C-III uses ubiquinol to reduce Cyt-c, which transfers the electrons to C-IV, which performs the reduction of O_2_ to H_2_O. The O_2_-producing activity of PSII provides enough O_2_ for respiration in the light. However, respiratory O_2_ consumption at low light or in darkness, under conditions of limited O_2_ exchange with the environment, can decrease O_2_ availability for PTOX in the chloroplast, resulting in a highly reduced PQ pool, which in turn results in the increase of Q_A_ reduction and an increased Chl fluorescence yield.

An important finding of our work is that the Chl fluorescence wave in intact corals can be induced not only by artificial removal of dissolved oxygen, but also by the respiratory activity of the host when equilibration of dissolved oxygen with a larger water reservoir in the vicinity of the corals is limited (Fig. 4D, E, and Scheme I). In situ measurements using sensitive nanoparticle-based oxygen sensors showed that the oxygen concentration inside the coral tissue in the dark can be as low as 35-65% of air saturation^38^. Such oxygen levels are low enough to alter the redox state of photosynthetic electron-transport components in the symbionts (Fig. 3A–E). This highlights how internal oxygen dynamics within coral tissues can modulate symbiont electron-transfer processes. Because dissolved oxygen availability in a symbiosome is lower than in the surrounding seawater, even relatively small environmental or physiological disturbances – such as reduced water flow^39^, moderate heat stress, the buildup of reactive oxygen species (ROS)^40,41^ caused by photodamage, or an elevated respiration rate in the host^42^ - can strongly affect symbiont physiology. Under such conditions, efficient metabolite exchange between host and symbiont becomes increasingly important for maintaining metabolic stability. Consequently, the observation of the fluorescence-wave phenomenon in both isolated symbionts and intact corals under stress indicates that it may be a useful, non-invasive tool for studying coral stress responses in situ.

An interesting feature of the slow-wave fluorescence phenomenon in intact corals is that the F_o_ fluorescence level shows a continuous increase and does not return to the initial baseline (e.g., Figs. 1-3) as happens in the isolated Symbiodiniaceae cultures (e.g., Figs. 3 and 6). The suppression of this dark F_o_ increase by DCMU (Fig. 7) confirms that the effect depends on the accumulation of reduced Q_A_, which is promoted by a more reduced plastoquinone (PQ) pool rather than by changes in intrinsic Chl fluorescence yield. This response is partly driven by oxygen consumption by the coral host (Figs. 4A–C), which lowers the availability of O₂ required for PTOX-mediated PQ oxidation. Reduced PTOX activity favors PQ reduction, which can be reinforced by stromal electron donors (e.g., NADPH) through chlororespiratory pathways (Scheme I). However, Q_A_ reduction in the dark, reported by the increase in F_o_ fluorescence, can also be observed when the glucose-glucose oxidase-catalase treatment lowers the dissolved O_2_ concentration near to anaerobic levels, below 1 μM/L, at which point O_2_ remains constant during the fluorescence measurements (Fig. 4 A and B). This shows that Q_A_ is gradually reduced in the host-embedded symbionts even when the O_2_ concentration is stable – an effect not observed in cultured symbionts (Fig. 3A). Thus, in addition to host respiration, another host-mediated process appears to contribute to Q_A_ reduction in intact corals. Such a process could be a direct or indirect electron transfer from the host to the symbiont, increasing the reduction state of the PQ pool, which is transmitted back to Q_A_ via charge-equilibration processes (see Scheme I and further discussion below).

### Extracellular electron transport revealed by Chl fluorescence relaxation characteristics

To obtain a deeper insight into the regulation of the redox state of the photosynthetic electron transport components in symbiont cells, we examined whether extracellular electron transfer processes might be involved. For this purpose, we used FeCN, a well-known external electron acceptor, which has been widely employed as an effective electron-transfer mediator for various plasma-membrane redox enzymes and transport protein^43,44^. Given that FeCN does not penetrate the plasma membrane of intact microalgal cells^23^, its reduction in cell suspensions is expected to occur primarily through electron-transfer processes directed toward the extracellular space. In *Fugacium* cells, the presence of FeCN resulted in a marked suppression of the middle phase of the fluorescence decay pattern and the elimination of the wave between10–400 s (Fig. 5). These effects are consistent with enhanced oxidation of the plastoquinone pool, potentially caused by FeCN acting as an external electron sink that draws electrons from plasma-membrane–associated redox pathways. This suggests that extracellular electron-transfer (EET) processes may contribute to the observed modulation of the photosynthetic redox state.

In intact coral nubbins, the FeCN-induced suppression of the F_o_ level, together with its retarded rise between 10-400 s (Figs. 6 and 7), indicates that the reduction level of Q_A_ is also decreased by FeCN in the host-embedded symbiont. However, the preserved middle phase of the fluorescence relaxation (Fig. 6B), as well as the continued increase of the F_o_ level after longer incubation times (Fig. 7), shows that Q_A_ (and the PQ pool) is kept more reduced *in hospite* than in isolated symbionts in the presence of FeCN. This effect is possibly caused by an electron transfer process, which is directed from the host to the symbiont. The detection of OJIP-type fluorescence induction traces (Fig. 8) confirm that the extracellular electron acceptor FeCN competes with the PQ pool for electrons in intact corals. The OJIP traces also show a slower fluorescence rise in the intact coral nubbins than in the isolated symbiont culture, which indicates that electrons can leak to the exterior of the symbiont cells *in hospite*, including possibly to the host as indicated in Scheme I.

### Extracellular electron transport revealed by ferricyanide reduction assay

The results of the FeCN assay in cultured *Fugacium* (Figs. 9-10) confirmed that extracellular electron transport takes place and FeCN functions as an alternative electron sink outside the cells. Interestingly, FeCN reduction proceeded at nearly the same rate in the dark and under illumination (Fig. 9), and DCMU caused only a minor suppression of this activity in *Fugacium*. This suggests that the majority of the electrons reducing FeCN do not originate directly from light-driven photosynthetic electron flow. The sustained FeCN reduction in the dark over periods of up to 15 hours—until all added FeCN was fully reduced—indicates that the electrons most likely originate from cellular metabolic reductants, such as NADH or NADPH, supplied by non-photosynthetic pathways. However, the effect of FeCN on the reduction level of Q_A_ indicates that these metabolic reductants affect the reduction level of the PQ – which in turn modulates Q_A_ reduction via charge equilibria – most likely via the chlororespiratory pathway (Scheme I). This observation differs from recent reports in other Symbiodiniaceae species, where DCMU treatment typically results in ∼90% inhibition of photocurrent^35^. This variation may reflect differences in physiological responses to environmental conditions, such as illumination quality, as the previous study employed red light^35^, whereas *Fugacium* in our experiments was tested under white light. In addition, species-specific traits, which are consistent with our earlier observations of distinct chlorophyll fluorescence patterns within the Symbiodiniaceae family^8^, can also be responsible for the different extents of the DCMU effect, highlighting the metabolic diversity among Symbiodiniaceae species.

**Figure 9:**
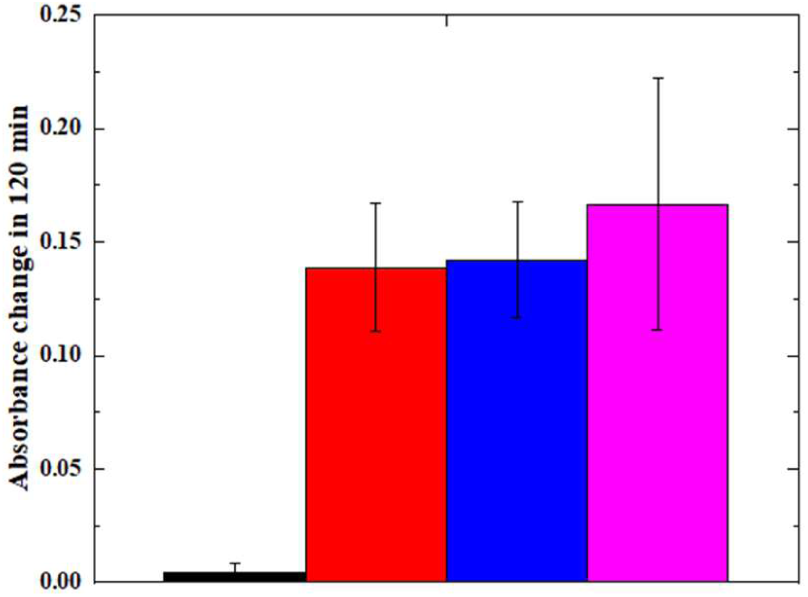
FeCN reduction in *Fugacium kawagutii* (CS156). Cells of 5μg Chl/mL were incubated in the presence of 1 mM FeCN in the dark (red) or under 200 μmol photons m^-2^s^-1^ illumination (blue) in the intact form, or after partial digestion of the cell walls (magenta). The black column shows the cell-free control. The absorbance change was detected at 420 nm after 2 hours. The data are mean values from 3 biological replicates

**Figure 10:**
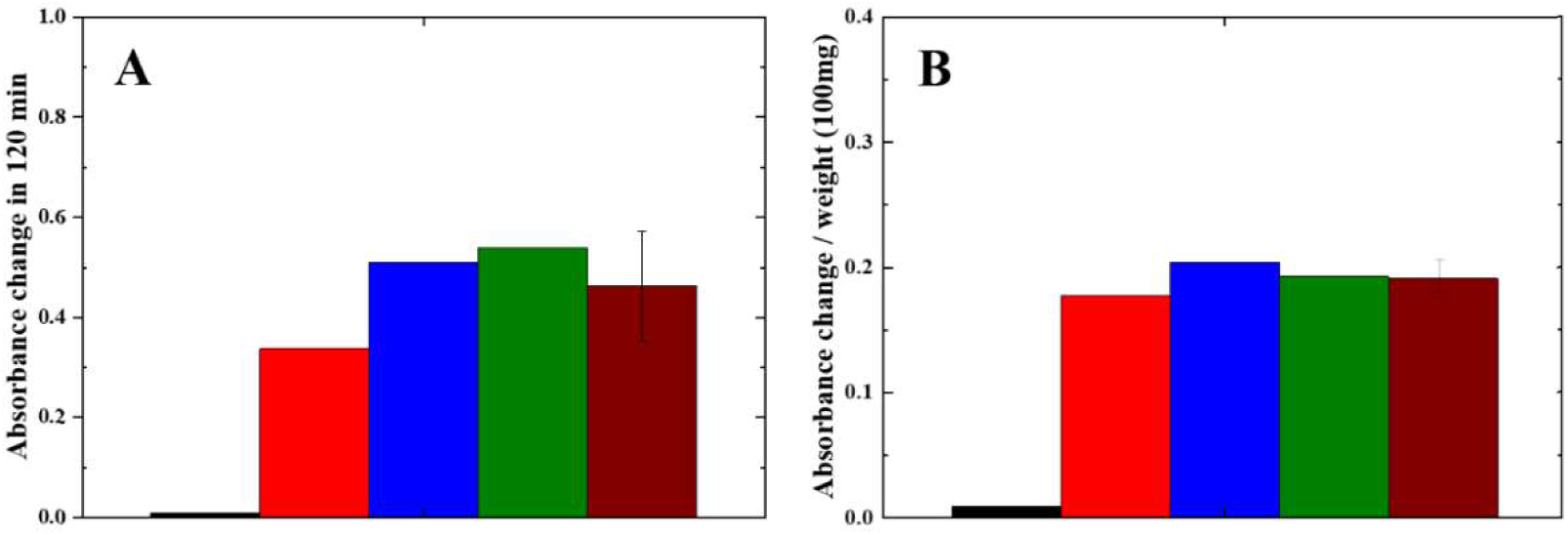
FeCN reduction by *Pocillopora damicornis.* (A) total absorbance change at 420 nm, (B) absorbance change per 100 mg of nubbin. The columns show the control without coral nubbin (black), individual coral nubbins (red, blue & green), and the average of the nubbins (brown).

The extracellular electron transfer activity of *Fugacium*, as quantified by the rate of FeCN reduction is 13.5 ± 2.4 µmol mg^-1^ Chl h^-1^, which outperforms cyanobacteria such as *Synechocystis* PCC6803, showing reduction rates ranging from 45 to 135 pmol nmol^-1^ Chl min^-1^ (3.0–9.1 µmol mg Chl^-1^ h^-1^) across various conditions, including high salinity, genetic modification, and their combination^23^. However, the EET activity of *Fugacium* is comparable to other eukaryotic microalgae such as *Chlorella vulgaris* (30.3 ± 0.8 µmol mg^-1^ Chl h^-1^), *Synechococcus elongatus* PCC7942 under iron-starved, neutral pH conditions (31 ± 1.0 µmol mg^-1^ Chl h^-1^), *Parachlorella kessleri* MACC-38 (∼38 µmol mg⁻¹ Chl h⁻¹), *P. kessleri* strain 27.87 (18 µmol mg⁻¹ Chl h⁻¹), and *P. kessleri* strains 211-11g and 211-11c (8–9 µmol mg⁻¹ Chl h⁻¹)^23,45^. Recently, Marcel et al. (2025) demonstrated using a bio-electrochemical system that *Symbiodinium* cells in culture exhibited an efficient EET, about 2.5 times higher than diatoms^35^.

With these EET rates, Symbiodiniaceae, together with other EET-capable microalgae, fall within the category of “poor” exoelectrogens, exhibiting limited but measurable extracellular electron transfer. The majority of electrons generated during light-driven reactions are consumed in CO₂ fixation and other intracellular processes, leaving only a limited number of electrons available from photosynthesis to be directed to extracellular electron transport (EET). The so-called “good” exoelectrogenic bacteria, such as *Shewanella oneidensis* and *Geobacter sulfurreducens,* do not require additional redox mediators and possess specialized adaptations that enable efficient coupling of respiration to extracellular electron acceptors. These bacteria employ outer membrane c-type cytochromes that facilitate direct electron transfer to external surfaces, in addition to redox-active molecules that facilitate electron flow outside the cell^22,46^. The main function of EET in *Shewanella* and similar bacteria is to reduce Fe(III) compounds, which cannot be taken up by the cells, to Fe(II), which can be biologically utilized. The growth and stress responses of Symbiodiniaceae also depend strongly on the availability of utilizable iron compounds^47–49^. However, the dominating form of iron in the seawater is Fe(III), which cannot be taken up directly by the cells. Although ligation of Fe(III) by organic compounds makes it available for cellular uptake, the reduction of Fe(III) to Fe(II) in the seawater can be an important function of EET in the Symbiodiniaceae cells.

In host-embedded symbionts, extracellular electron transport from the symbiont could occur not only to external acceptors located outside the coral but, in principle, also to the host, if appropriate external electron acceptors are available (see Scheme I). The observed decrease in the reduction level of Q_A_ in the presence of FeCN indicates that extracellular oxidants can influence the photosynthetic redox poise, most likely by drawing electrons from cellular metabolic pools that are ultimately in redox communication with the PQ pool. Even though riboflavin is a known EET mediator^50^, and its transporter is present in the outer membrane of both the symbiont and the host^29^, riboflavin - with its low redox potential (ca. −199 to −290 mV)^51–53^ is unlikely to act as an electron shuttle from the symbiont to the host, since it cannot accept electrons energetically downhill from the PQ pool (Eₘ ≈ +110 mV)^54^. If EET occurs, it is more plausible that plasma-membrane–associated oxidoreductases or membrane-bound redox proteins mediate electron release, although specific pathways remain to be identified.

Regarding potential candidates for accepting electrons in the host from the symbiont, high redox-potential components in the mitochondrial respiratory electron transport chain could be considered. Given the relatively high redox potential of the ferricyanide/ferrocyanide couple (≈ +360 mV), which acts as an efficient EET acceptor in both isolated symbionts and intact corals (Figs. 9, 10), host electron acceptors would likely need to have comparably high potentials. For example, cyt-c in the mitochondrial electron transport chain (E_m_: +220 to + 270 mV)^55,56^, could potentially fulfill this role, although its redox potential is somewhat lower than that of FeCN. It should be noted, however, that currently we do not have direct evidence that the electrons reducing FeCN outside the symbiont cells are actually accepted by host electron transport components, aside from the slower rise of the OJIP kinetics in the coral nubbins compared to the isolated symbiont cultures.

The gradual increase of the baseline fluorescence under very low dissolved oxygen levels (Fig. 4B), indicative of Q_A_ reduction in the host-embedded symbionts even under minimal host respiration, raises the intriguing possibility that electrons can be transferred not only from the symbiont to external acceptors, but also in the opposite direction - from the host to the symbiont. Uptake of electrons from extracellular sources (EEU) is a known phenomenon in anaerobic microbes, which are capable of EET^31,32^, and can exchange electrons bidirectionally with their environment. Potential sources of electrons for the putative host-to-symbiont transfer could include low-potential cellular reductants. For example, FADH₂ in the mitochondrial electron transport chain could, in principle, contribute electrons if converted to a diffusible mediator such as riboflavin, which is an efficient EET shuttle^50^ and has transporters in both host and symbiont membranes^29^. Therefore, riboflavin could potentially act as an electron shuttle between the host and the symbiont. Other candidates, such as ubiquinone (denoted as CoQ in Scheme I), are membrane-bound –like PQ in the symbiont - could also contribute to redox interactions in principle. The lower redox potential of riboflavin (below −200 mV) and of CoQ (ca. 0 mV)^51–53^ relative to the PQH_2_/PQ in the symbiont (at around +110 mV)^54^, would make such an electron transport energetically feasible, although the precise pathways remain speculative.

Our working hypothesis is that besides electron transport from the symbiont to external acceptor located outside the coral, such as Fe(III) in the seawater, there could be a bidirectional electron exchange between the zooxanthellae and coral host cells. The largely light-independent reduction rate of FeCN shows that EET can occur in the dark, when sufficient extracellular electron acceptors are present. The source of electrons for this process is most likely chlororespiration, which delivers electrons to the PQ pool from NADPH via the NDH-2 complex. The electrons putatively transferred from the symbiont to the host could enhance the efficiency of respiration in the mitochondria. The reverse process - extracellular electron uptake from the host by the symbiont - could, in principle, occur under low oxygen conditions or in the dark when low-potential components of the host respiratory electron transport chain, such as FAD and CoQ, are in a reduced state due to limited O₂ availability. This bidirectional electron exchange likely plays an important role in maintaining coral health, particularly under stress, by enabling efficient redox balancing and demonstrating how mutualistic relationships can enhance metabolic resilience.

## Conclusions

While metabolic exchange between the symbiont and host, including nutrient transfer from host to symbionts, is well documented^29,34,58^, direct evidence for electron-level interactions has not been reported. Our results reveal a complex interplay between photosynthetic and host respiratory electron transport in coral symbiosis. Four key conclusions emerge:

1. **Host respiration indirectly affects photosynthetic electron transport**, as evidenced by Q_A_ reduction in the symbiont via mitochondrial oxygen consumption.
2. **Symbionts can transfer electrons extracellularly**, both as isolated cells and *in hospite*, to the artificial acceptor FeCN. This Fe(III)-to-Fe(II) reduction suggests that extracellular electron transport could similarly reduce Fe(III) to Fe(II) in seawater, facilitating uptake of bioavailable iron.
3. **Electron transfer from symbiont to host is indicated** by slower OJIP fluorescence kinetics in intact corals compared to isolated Symbiodiniaceae, suggesting PQ pool electrons may be drained toward the host.
4. **Electron transfer from host to symbiont is also possible**, as shown by sustained Q_A_ reduction in intact corals—but not in cultures—under microaerobic conditions when host-respiration-induced Q_A_ reduction does not occur.

Previous studies have shown correlations between host respiration and symbiont photosynthesis^57,59^, and the coupling of these processes may benefit metabolic exchange^60^. Our findings further suggest that this interaction involves previously unrecognized extracellular electron exchange pathways between host and symbiont. Future work is needed to identify the specific mechanisms mediating electron flow in both directions.

## Materials and methods

### Conditions for Symbiodiniaceae cultures and the coral tank

*Fugacium kawagutii* (CS156, formerly clade F1), was cultivated at 24°C under 50 µmol photons m^-2^ s^-1^ white light (12h:12h light: dark) in F/2 media for one week, reaching the mid-log growth phase. Cells were centrifuged at 5000× g for 13 min and re-suspended in fresh F/2 media. The final chlorophyll (Chl) concentration of the cultures was determined based on extraction with 100% methanol^61^. The Chl content was adjusted (using fresh F/2 medium) to 5 μg/mL for the experimental samples. Prepared samples were maintained under growth light and temperature conditions for 1 h to acclimate before each measurement.

For the coral tanks, small nubbins of *Pocillopora damicornis*, *Acropora* and *Seriatopora* were glued to rocks and cultivated within a controlled tank environment with water parameters maintained as: Calcium (425-440 mg/L), Magnesium (1350-1380 mg/L), Alkalinity (7.8-8 KH), Phosphate (0.03-0.05 mg/L), Nitrate (3-5 mg/L), and Specific Gravity (1.025-1.026), which were monitored every second day. The pH exhibited fluctuations, ranging from 8.1 at night to 8.35 during the day. Each week, 10% of the water was refreshed with Red Sea salt water with a water flow set to 33 times turnover per hour, and lighting maintained at a PAR level of 250-350 µmol photons m^-2^ s^-1^.

### Measurement of variable Chl chlorophyll fluorescence relaxation

Chlorophyll *a* fluorescence induction curves (OJIP transients) were measured using a double-modulation fluorimeter (FL-3000, Photon Systems Instruments, Drásov, Czech Republic). The intensity of the saturation pulse was set to 3500 μmol m^-2^ s^-1^ with a duration of 3 seconds. The wavelength of the saturation pulse was 650 nm.

The measurement of flash-induced chlorophyll fluorescence relaxation kinetics (FF) was conducted using a double-modulation fluorimeter (FL-3000, Photon Systems Instruments, Drásov, Czech Republic)^62^. Algae samples and coral nubbins were placed in a cuvette with a 1 cm path length and dark adapted for 3 min before each measurement.

To initiate the measurement, four measuring flashes with a duration of 8 μs, separated by intervals of 200 μs (wavelength of 620 nm), were applied to determine the minimum fluorescence in the dark (F_o_). Subsequently, a single turnover saturating actinic flash (30 μs, wavelength of 639 nm) was administered to induce the formation of Q_A_^-^, resulting in a rise in fluorescence intensity. This maximal fluorescence recorded after the single-turnover actinic flash was specifically denoted as F_m(ST)_. The relaxation of flash-induced fluorescence yield increase typically exhibits a monotonous decrease which can be assigned into three distinct phases: (i) a fast decay phase with a duration of approximately 300–500 μs due to Q_A_^-^ reoxidation by Q_B_ (or Q_B_^-^), (ii) a middle phase due to reduction of plastoquinone (PQ), which binds to the Q_B_ site after the flash, lasting about 5–15 ms, and (iii) a slow phase as a result of recombination of the electron on Q_A_Q_B_^-^ with the oxidized S_2_ (or S_3_) state of the water-oxidizing complex, which lasts approximately 10–20 s^17^. The measuring flashes were recorded in the time range from 150 μs to 400 s on a logarithmic time scale, to study the fluorescence relaxation at a much longer time scale.

### Dissolved oxygen measurements

Oxygen evolution and uptake rates were measured using two different electrode systems: the Fibox3 oxygen sensor (PreSens Precision Sensing GmbH, Germany) and the DW2 electrode chamber equipped with an S1 Clark-type O_2_ electrode (Hansatech Instruments Ltd, UK). For measurements using the PreSens Fibox3 electrode, the system was first calibrated using 100 mL of bubbled distilled water (representing the dissolved oxygen content of ambient air) and 100 mL of a solution containing 1 g of sodium sulfite (Na_2_SO_3_) (representing the anoxic condition). Following calibration, the electrode was placed in the cuvette, and the decrease in dissolved oxygen was measured along with fluorescence relaxation kinetics.

In the case of using the Clark-type polarographic oxygen electrode, two-point calibration was performed, at ambient air dissolved oxygen concentration and at oxygen-depleted condition (medium was flushed with N_2_ gas). 1 mL of *Fugacium* culture (5 μg mL^-1^ Chl) or a coral nubbin in 1 mL of tank media was placed in the reaction chamber, dark-adapted for 3 minutes at 24°C under constant stirring. Oxygen evolution was measured through a 2 min dark phase followed by a 2 min light phase (200 μmol photons m^-2^ s^-1^) with the help of a fiber optic light source Schott KL 1500 (Schott AG) connected to one side. The light-dependent oxygen evolution rate was calculated as the difference between oxygen consumption in the dark and oxygen production in the light. This procedure was also carried out with intact coral nubbins, which were subsequently used for the ferricyanide (FeCN) reduction assay. The chlorophyll concentration in the nubbins was later normalized by comparing the oxygen evolution rate of each nubbin with the rates measured for *Fugacium* cultures containing 5 μg mL^-1^ Chl.

Microaerobic conditions during measurements with *Fugacium* cultures or intact coral nubbins were established enzymatically by supplementing the medium with 10 mM glucose, 7 U ml^-1^ glucose oxidase, and 60 U ml^-1^ catalase, followed by dark incubation for 15 minutes. Flash-induced Chl fluorescence relaxation curves were subsequently measured under these microaerobic conditions.

### Ferricyanide (FeCN) reduction assay

For the investigation of extracellular electron transfer, FeCN reduction assay was performed with 1 mM concentration. *Fugacium* cells and coral nubbins were incubated in 1 mM FeCN solution on a shaker under 200 µmol photons m^-2^ s^-1^ light intensity. Absorption spectrum of the filtrate of the FeCN solution incubated with *Fugacium* cells or coral nubbins (1 mL filtrate was obtained using a GF/C glass fibre filter, Whatman) was recorded between 380 and 650 nm using UV-1800 spectrophotometer (Shimadzu Corporation, Japan). The change at 420 nm over a period of 120 min was observed. The rate of reduction was calculated with the formula^23^.

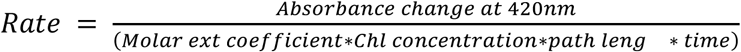

### Experimental procedure

Flash-induced chlorophyll fluorescence relaxation kinetics (FF) was recorded on different species of corals in *Fugacium* cell cultures. The 1-400 second long relaxation phase was analyzed to reveal changes in fluorescence that are related to the slow electron transport processes leading to the reduction of PQ pool. The Photosystem II electron transport inhibitor DCMU, and the extracellular electron acceptor potassium ferricyanide (FeCN) was applied at 20 µM and 1 mM concentration, respectively, when indicated in the figures. Experiments were also performed with and without actinic flash, and with different oxygen conditions measured simultaneously along with PreSens electrode on intact coral. FeCN reduction assays were conducted on isolated *Fugacium* cultures, either in the intact state or after partially digesting the cell walls, and intact coral nubbins. To allow comparison of *Fugacium* cell cultures and coral nubbins the FeCN reduction rates were normalized to equal Chl contents by using the assumption that O_2_-evolving capacity of *Fugacium* cells is identical in isolated cell cultures and *in hospite*.

## Acknowledgements

M.S. was funded by the National Research, Development and Innovation Office (NKFIH) grant FK 146298. Sz.K. was funded by the National Research, Development and Innovation Office (NKFIH) grant PD 146655

## Author contributions

I.V. conceived the idea, designed and interpreted the experiments, and finalized the manuscript. S.M.A. performed the majority of the experiments, prepared the figures, and wrote the initial version of the manuscript. P.P.P performed part of the Chl fluorescence relaxation measurements in isolated *Fugacium* cells. M.S. supervised the experimental work, participated in the interpretation of the results, writing, correcting, and finalizing the manuscript. Sz.K. took care of growing and preparing the coral samples. A.W.D.L participated in the initial experiments in the Heron Island Research Station, as well as in finalizing the manuscript.

